# Marine fish intestine responds to ocean acidification producing more carbonate aggregates

**DOI:** 10.1101/119388

**Authors:** Sílvia F. Gregório, Ignacio Ruiz-Jarabo, Edison M Carvalho, Juan Fuentes

## Abstract

Marine fish contribute to the carbon cycle by producing mineralized intestinal aggregates generated as by-products of their osmoregulation. Here we aimed at characterizing the control of intestinal aggregate production in the gilthead sea bream in response to near future increases of environmental CO_2_. Our results demonstrate that hypercapnia (800 and 1200 μatm CO_2_) elicits higher intestine epithelial HCO_3_^−^ secretion and the subsequent parallel increase of intestinal aggregate production when compared to present values (400 μatm CO_2_). Intestinal gene expression analysis revealed the up-regulation of crucial transport mechanisms involved not only in the intestinal secretion cascade (*Slc4a4, Slc26a3 and Slc26a6*) of sea bream, but also in other mechanisms involved in intestinal ion uptake linked to water absorption such as *NKCC2* and the *Aquaporin 1b.* These results highlight the important role of fish in the marine carbon cycle, and their potential growing impact of intestinal biomineralization processes in the scenario of ocean acidification.

## INTRODUCTION

In seawater, fish live in a hyperosmotic environment which leads to a loss of body fluids. The water lost passively through the epithelial surface is compensated by drinking (Fuentes and Eddy, 1997; Smith et al., 1930). Due to the osmotic imbalance between the ingested SW and the internal body fluids, an intense desalinization in the first part of the digestive tract is required i. e. oesophagus and stomach (Hirano and Mayer-Gostan, 1976; Parmelee and Renfro, 1983). This ion capture, mainly Na^+^ and Cl^−^, results in the decrease of intestinal fluid osmolality from circa 1050 to 400 mOsm kg^−1^ due to the action of active and passive transport processes (Esbaugh and Grosell, 2014; Loretz, 1995; Marquez and Fuentes, 2014). The osmotic pressure of the intestinal fluid is then decreased to a level that matches plasma osmolality (Gregorio et al., 2013), allowing the water to be absorbed passively or by the action of aquaporins (Whittamore, 2012; Wood and Grosell, 2012).

Water absorption in the anterior part of the intestine of marine fish was associated to ion fluxes driven by Na^+^/K^+^/2Cl^−^ co-transporters (Musch et al., 1982). However, it was recently established that part of the process is also linked to the action of chloride-dependent movements coupled to bicarbonate secretion, and mediated by apical anion transporters (Grosell et al., 2005; Heuer et al., 2012). Members of the Slc26 family are responsible for this apical mechanism in fish enterocytes (Carvalho et al., 2015; Gregorio et al., 2013; Ruhr et al., 2016). HCO_3_^−^ enters the cell through a basolateral Na^+^/HCO_3_^−^ co-transporter belonging to the Slc4 family (Chang et al., 2012) or is formed by the intracellular action of carbonic anhydrases that hydrate the CO_2_ resulting in the production of HCO_3_^−^ and H^+^ ions (Grosell et al., 2009).

The secretion of HCO_3_^−^ into the intestinal luminal fluid promotes an increase of pH (Wilson and Grosell, 2003). In parallel calcium and magnesium are present in high concentrations in SW and accumulate in the intestinal fluid. Under both conditions, precipitation and formation of carbonate aggregates in the intestinal lumen of marine fish is promoted. As a result of this process, an osmolality reduction of the fluid occurs, thus facilitating water absorption (Wilson et al., 2009; Wilson et al., 2002). Apical bicarbonate secretion in the enterocyte is regulated by biotic factors such as hormones, peptides, blood HCO_3_^−^ concentration and abiotic factors *e.g.* salinity (Genz and Grosell, 2011; Genz et al., 2008; Gregorio et al., 2013). On the other hand, previous studies demonstrated that bicarbonate secretion in the intestine of marine fish is also regulated by an intracellular pH/CO_2_/HCO_3_^−^ sensor, the soluble adenylyl cyclase (Carvalho et al., 2012; Tresguerres et al., 2010). Moreover, a challenge with high environmental levels of CO_2_ (hypercapnia) elicited a response in the gilthead sea bream (*Sparus aurata*), decreasing its plasma pH during the innitial 24 h, which is then buffered by increased levels of plasma bicarbonate (Michaelidis et al., 2007). Such an accumulation of bicarbonate in plasma, but also in other body fluids was also observed in several organisms (Portner et al., 2004). It should be of interest to establish if those physiological modifications alter intestinal carbonate aggregates production in marine fish.

In a global scenario, as a result of anthropogenic emissions in the atmosphere CO_2_ increased from 280 up to 380 μatm in the 20^th^ century. The ocean absorbs part of this CO_2_ resulting in a decrease of pH (IPCC, 2007; Kroeker et al., 2013). The ocean surface CO_2_ levels are predicted to duplicate by the year 2100 to achieve levels circa 800 μatm CO2, thus decreasing 0.3-0.4 pH units (Orr et al., 2005). Previous studies have reported that some marine calcifying biota is affected when submitted to elevated CO_2_ levels and the subsequent pH decrease close to values expected by the year 2100 (pH 7.8) and by the end of the next century (pH 7.6) (Mackenzie et al., 2014; Munday et al., 2009; Munday et al., 2010). Moreover, by the year 2300 the seawater CO_2_ concentration will be 1900 μatm that in turn decreased the pH in up to 0.77 units (Caldeira and Wickett, 2003; Kroeker et al., 2013; Zeebe et al., 2008). In contrast to other taxa, marine fish contribute to the carbon cycle as they fix/cycle around 15 *%* of the oceanic carbon, producing mineralized intestinal aggregates generated as by-products of their osmoregulation (Walsh et al., 1991; Wilson et al., 2009). The nature of those aggregates makes them relevant as they provide neutralizing alkaline buffer in the form of soluble carbonates, preferentially in the uppermost, most productive section of the water column (Perry et al., 2011). This study aimed at understanding the intestinal response and control of aggregate production of marine fish intestine in response to near future concentrations of CO2 in seawater.

## METHODS

### Animals

Sea bream (*Sparus aurata*) juveniles were purchased from CUPIMAR SA (Cadiz, Spain) and transported to Ramalhete Marine Station (CCMAR, University of Algarve, Faro, Portugal). Fish were maintained for 60 days in 1000 L tanks with running seawater at a density 9-10 kg m^−3^ and fed 2 *%* ration (fish wet weight, Sorgal, S.A., Portugal; Balance 3) twice daily until the start of the experiment. For experimental purposes 5 fish (200 g body weight) were transferred to 100 L tanks. Eight replicates were established per CO_2_ concentration in an open circuit were temperature was maintained constant (25 °C), photoperiod was natural (April-June, Algarve, Portugal) and the feeding regime was maintained as above. Food was withheld for 36 h before sacrifice and sampling. No mortality was observed during the experiments.

The experiments comply with the guidelines of the European Union Council (86/609/EU). All fish protocols were performed under a ‘‘Group-C’’ licence from the Direcção-Geral de Veterinária, Ministério da Agricultura, do Desenvolvimento Rural e das Pescas, Portugal.

### Experimental conditions and seawater chemistry

The rate of CO_2_ injection into the systems was controlled by the pH level of seawater using pH probes connected to CO_2_ controllers (EXAxt PH450G; Yokogawa Iberia-Portugal) with an internal controller following guidelines provided by the manufacturer. Each independent circuit was injected with CO_2_, thus obtaining three groups constantly maintained at 400, 800 and 1200 μatm CO_2_. To calibrate the automated negative feedback system, it was necessary to measure seawater pH, and extrapolate a pH/*p*CO_2_ standard curve to indirectly measure *p*CO_2_. The linearity of the relationship was confirmed through IRGA CO_2_ analysis (WMA-4 PP Systems, Amesbury, MA, USA).

Total alkalinity (TA) was measured using an automated, open-cell potentiometric titration unit (SOP 3b) (Dickson et al., 2007), with a combination DL15 titrator and a DG115-SC probe (Mettler-Toledo) using certified acid titrant (0.1 M HCl, Fluka Analytical, Sigma-Aldrich). The results are expressed as μmol kg^−1^ seawater. The water chemistry parameters were calculated using the CO_2_Calc Software (Robbins et al., 2010). Along with temperature and salinity, CO_2_Calc algorithms produce a suite of carbonate system parameters using only two measured variables (of five potential inputs); the remaining three possible inputs are also calculated as algorithm outputs. In our samples, pH, TA, and/or TCO_2_ were routinely measured, and precedence of use is: 1) measured pH and TA 2) measured pH and TCO_2_ (total dissolved inorganic carbon). In situ carbonate system parameters were also calculated using temperature recorded in the field (“adjusted” outputs), as explained before (Robbins et al., 2010).

### General sampling

After 3 months of acclimation to the altered water CO_2_ levels, fish were captured and anesthetized in 2-phenoxyethanol (1: 10,000 v/v; Sigma-Aldrich, St Louis, MO, USA). Blood samples were collected by caudal puncture using heparinized syringes. Plasma was obtained by centrifugation (10,000 *rpm,* 5 min, 4 °C) and stored at −20 °C until analysis. Fish were sacrificed by decapitation and the whole intestine was isolated. The intestinal fluid of individual fish was collected from the excised intestinal tract clamped (from pyloric caeca to anal sphincter) with two haemostatic forceps, emptied into pre-weighed vials and centrifuged (12,000 *rpm,* 5 min, RT) to separate fluid from precipitates. The fluid was transferred to pre-weighed vials and the volume was measured to the nearest 0.1 μL (0.1 mg, assuming a density of 1).

### Plasma and fluid analysis

Osmolality of plasma and intestinal luminal fluid was measured in 10 μL samples with a Vapro 5520 osmometer (Wescor, South Logan, UT, USA). Sodium concentrations were measured by flame photometry (BWB-XP Performance Plus, BWB Technologies, UK). The results are expressed as mmol L^−1^. Chloride was determined by Coulomb-metric titration (SAT-500, DKK-TOA, Japan). Calcium and magnesium were measured by colorimetric tests, using commercial kits (Spinreact, Reactivos Spinreact, SA, Girona, Spain), according to the manufacturer instruction in a microplate reader (Biorad Benchmark, Bio-Rad, USA). Intestinal fluid titratable alkalinity (HCO_3_^−^ + CO_3_^2−^) was manually measured with the double titration method with a pH electrode (HI1330B, Hanna Instruments, Smithfield, RI, USA) attached to a pH meter (PHM84, Radiometer, Copenhagen, Denmark): 50 μL of the intestinal fluid samples was diluted in 10 mL NaCl (40 mmol L^−1^), gassed with CO_2_-free gas for 30 min to remove CO_2_ and titrated to pH 3.8 with 10 mmol L^−1^ HCl; an additional gassing period of 20 min was applied to remove any remaining CO_2_. The sample was back titrated to its original pH with 10 mmol L^−1^ NaOH. The volume difference between added acid and base in both titrations and titrant molarities was used to calculate total HCO_3_^−^ equivalents (mEquiv.L^−1^) as described before (Gregorio et al., 2014).

### Intestinal Bicarbonate Secretion

A segment of fish anterior intestine was excised, mounted on tissue holders (P2413, 0.71 cm^2^, Physiologic Instruments, San Diego, CA, USA) and positioned between two half-chambers (P2400, Physiologic Instruments) containing 1.5 mL of basolateral and apical saline. The composition of the basolateral saline was: 160 mmol L^−1^ NaCl, 1 mmol L^−1^ MgSO_4_, 2 mmol L^−1^ NaH_2_PO_4_, 1.5 mmol L^−1^ CaCl_2_, 5 mmol L^−1^ NaHCO_3_, 3 mmol L^−1^ KCl, 5.5 mmol L^−1^ glucose and 5 mmol L^−1^ HEPES, pH 7.800, gassed with 0.3% CO_2_+ 99.7% O_2_. Apical saline: 88 mmol L^−1^ NaCl, 9.5 mmol L^−1^ MgCl_2_, 3 mmol L^−1^ KCl, 7.5 mmol L^−1^ CaC_2_, 126.5 mmol L^−1^ MgSO_4_ and 1 mmol L^−1^ Na_2_HPO_4_, gassed with 100% O_2_ and pH maintained at 7.800 throughout the experiments by pH-Stat (see below). The temperature was maintained at 25 °C throughout all experiments. All bioelectrical variables were monitored by means of Ag/AgCl electrodes (with tip asymmetry <1 mV) connected to either side of the Ussing chamber with 3 mm- bore agar bridges (1 mol L^−1^ KCl in 3% agar). Transepithelial electrical potential (TEP, mV) was monitored by clamping of epithelia to 0 μA cm^−2^. Epithelial resistance (Rt, Ω cm^2^) was manually calculated (Ohm’s law) using the voltage deflections induced by a 10 μA cm^−2^ bilateral pulse of 2 s every minute. Current injections were performed by means of a VCC 600 amplifiers (Physiologic Instruments). For pH-Stat control, a pH electrode (PHC 4000-8, Radiometer) and a microburette tip were immersed in the luminal saline and connected to a pH-Stat system (TIM 854, Radiometer). To allow pulsing (for Rt calculation) during pH measurements, the amplifier was grounded to the titration unit. The configuration of amplifier/pH-Stat system used in this study is similar to that first established for the characterization of HCO_3_^−^ secretion in the intestine of the Gulf toadfish and sea bream (Gregorio et al., 2013; Grosell and Genz, 2006; Guffey et al., 2011; Taylor et al., 2010) and provides rates of intestinal secretion similar to those obtained by the double titration method in the intestine of the sea bream (Fuentes et al., 2010). Measurement of HCO_3_^−^ secretion was performed on luminal saline at physiological pH 7.800 during all experiments. The volume of the acid titrant (2.5 mmol L^−1^ HCl) was recorded and the amount of HCO_3_^−^ secreted (nmol h^−1^ cm^−2^) was calculated from the volume of titrant added, the concentration of the titrant and the surface area (cm^2^). All experiments comprised 1 h of tissue stable voltage and HCO_3_^−^ secretion.

### qPCR

After anaesthesia and decapitation, a portion of the anterior intestine was collected from individual fish, and stored in RNA Later at 4 °C (Sigma-Aldrich) until utilized for RNA extraction within 2 weeks. Total RNA was extracted from samples of anterior intestine with the Total RNA Kit I (E.Z.N.A, Omega, US) following the manufacturer’s instructions and the quantity and quality of RNA assessed (Nanodrop 1000, Thermo Scientific, US). Prior to cDNA synthesis RNA was treated with DNase using the DNA-free Kit (Ambion, UK) following the supplier’s instructions. Reverse transcription of RNA into cDNA was carried out using the RevertAid First Strand cDNA Synthesis Kit (TermoFisher Scientific, UK) with 500 ng of total RNA in a reaction volume of 20 μL. Table 4 shows primer sequences and amplicon sizes.

Real-time qPCR amplifications were performed in duplicate in a final volume of 10 μL with 5 μL SsoFast EvaGreen Supermix (Bio-Rad, UK) as the reporter dye, around 20 ng cDNA, and 0.3 μM of each forward and reverse primers. Amplifications were performed in 96-well plates using the *One-step Plus* sequence detection system (Applied Biosystems, California, USA) with the following protocol: denaturation and enzyme activation step at 95 °C for 2 min, followed by 40 cycles. After the amplification phase, a temperature-determining dissociation step was carried out at 65 °C for 15 s, and 95 °C for 15 s. For normalization of cDNA loading, all samples were run in parallel using 18S ribosomal RNA (*18S*). To estimate efficiencies, a standard curve was generated for each primer pair from 10-fold serial dilutions (from 0.1 to 1e10^−8^ ng) of a pool of first-stranded cDNA template from all samples. Standard curves represented the cycle threshold value as a function of the logarithm of the number of copies generated, defined arbitrarily as one copy for the most diluted standard. All calibration curves exhibited correlation coefficients R^2^>0.98, and the corresponding real-time PCR efficiencies between 95 and 99%.

### Statistics

All results are shown as mean ± standard error of the mean (mean ± s.e.m.). After assessing homogeneity of variance and normality, statistical analysis of the data was carried out by using one-way analysis of variance using CO_2_ concentration as a factor of variance, followed by the *post hoc* Bonferroni t-test (Prism 5.0, GraphPad Software for McIntosh). The level of significance was set at p< 0.05.

## RESULTS

No mortality was observed in any of the experimental groups.

### Experimental conditions - seawater chemistry

The environmental condition values were controlled twice a day in this study and the results are shown in Table 1. Temperature and salinity levels were maintained within a small range of variation, and no differences were described between CO_2_ treatments (Table 1). The concentration of CO_2_ and pH in the control treatment averaged 408 ± 20 μatm and pH 8.01 ± 0.01, in the medium CO_2_–concentration group 797 ± 29 μatm and pH 7.80 ± 0.01, whereas in the enriched-CO_2_ treatment it averaged 1259 ± 46 μatm and pH 7.64 ± 0.01. Total alkalinity was not significantly different between treatments ranging between 2582 and 2605 μmol kg^−1^ SW (Table 1).

**Table 1.** Chemical conditions of seawater where sea bream were kept under different CO_2_ concentrations over a three month period.

| | 400 μatm CO<sub>2</sub> | 800 μatm CO<sub>2</sub> | 1200 μatm CO<sub>2</sub> |
| --- | --- | --- | --- |
| pH | 8.01 ± 0.01 | 7.80 ± 0.01 | 7.64 ± 0.01 |
| pCO<sub>2</sub> (μatm) | 408 ± 20 | 797 ± 29 | 1259 ± 46 |
| Salinity (ppt) | 37.5 ± 0.0 | 37.5 ± 0.0 | 37.5 ± 0.0 |
| Alkalinity (μMol kg<sup>-1</sup> SW) | 2605 ± 10 | 2582 ± 25 | 2585 ± 14 |
| T (°C) | 25.4 ± 0.2 | 25.5 ± 0.2 | 25.6 ± 0.2 |

### Plasma and intestinal fluid analysis

The osmolality and the ion composition of plasma and intestinal fluid of sea bream are shown in Table 2 and 3, respectively. No differences were detected in plasma osmolality which ranged between 343 to 346 mOsm kg^−1^ in controls and fish exposed to elevated CO_2_ concentrations (Table 2). Additionally, no changes in Cl^−^ (163.5 to 165.3 mmol L^−1^) and Na^+^ (178.7 to 186.2 mmol L^−1^) concentration (Table 2) were observed in the plasma between controls and fish exposed to elevated water CO_2_. The content of Ca^2+^ (9.5 ± 0.6 mmol L^−1^) in intestinal fluid did not change with the different exposure to elevated water CO_2_ (Table 3) and the concentration of Na^+^ (72.1 to 81.8 mmol L^−1^) in the intestinal fluid was not different between controls and fish exposed to elevated CO_2_ (Table 3). The content of Cl^−^ in the intestinal fluid of fish exposed to both levels of elevated water CO_2_ was significantly lower 95.0 ± 5.3 mmol L^−1^ (*p*<0.05, one-way ANOVA) when compared to controls 116 ± 5.5 mmol L^−1^ (Figure 1A). However, the content of Mg^2+^ in the intestinal fluid of fish exposed to elevated water CO_2_ was statistically higher than in control fish (*p*<0.05, one-way ANOVA, Table 3).

**Table 2.** Osmolality, Na^+^ and Cl^−^ levels in plasma of sea bream acclimated for 3 months to different CO_2_ concentration (400, 800 and 1200 μatm) at 37 ppt. Values are means ± s.e.m. (N=11-19). No differences were described between any of the experimental groups (p>0.05, one-way ANOVA).

| | 400 μatm CO<sub>2</sub> | 800 μatm CO<sub>2</sub> | 1200 μatm CO<sub>2</sub> |
| --- | --- | --- | --- |
| Osmolality (mOsm kg<sup>-1</sup>) | 342 ± 2 | 343 ± 2 | 346 ± 2 |
| Na<sup>+</sup> (mmol L<sup>-1</sup>) | 178.7 ± 2.3 | 183.4 ± 2.6 | 186.2 ± 2.1 |
| Cl<sup>-</sup> (mmol L<sup>-1</sup>) | 163.5 ± 1.3 | 165.2 ± 1.1 | 164.6 ± 1.3 |

**Table 3.** Osmolality, Na^+^, Ca^2+^ and Mg^2+^ levels in intestinal fluid of sea bream acclimated for 3 months to different CO_2_ concentration (400, 800 and 1200 μatm) at 37 ppt. Values are means ± s.e.m. (N=8-18). Different letters indicate significant differences among groups (*p*<0.05, one-way ANOVA).

| | 400 μatm CO<sub>2</sub> | 800 μatm CO<sub>2</sub> | 1200 μatm CO<sub>2</sub> |
| --- | --- | --- | --- |
| Osmolality (mOsm kg<sup>-1</sup>) | 353 ± 8 | 355 ± 7 | 359 ± 9 |
| Na<sup>+</sup> (mmol L<sup>-1</sup>) | 77.0 ± 2.4 | 72.1 ± 4.3 | 81.8 ± 5.0 |
| Ca<sup>2+</sup> (mmol L<sup>-1</sup>) | 9.5 ± 0.6 | 10.3 ± 0.6 | 8.6 ± 0.5 |
| Mg<sup>2+</sup> (mmol L<sup>-1</sup>) | 123.2 ± 4.3 <sup>a</sup> | 143.9 ± 7.8 <sup>b</sup> | 143.8 ± 6.5 <sup>b</sup> |

**Table 4.**
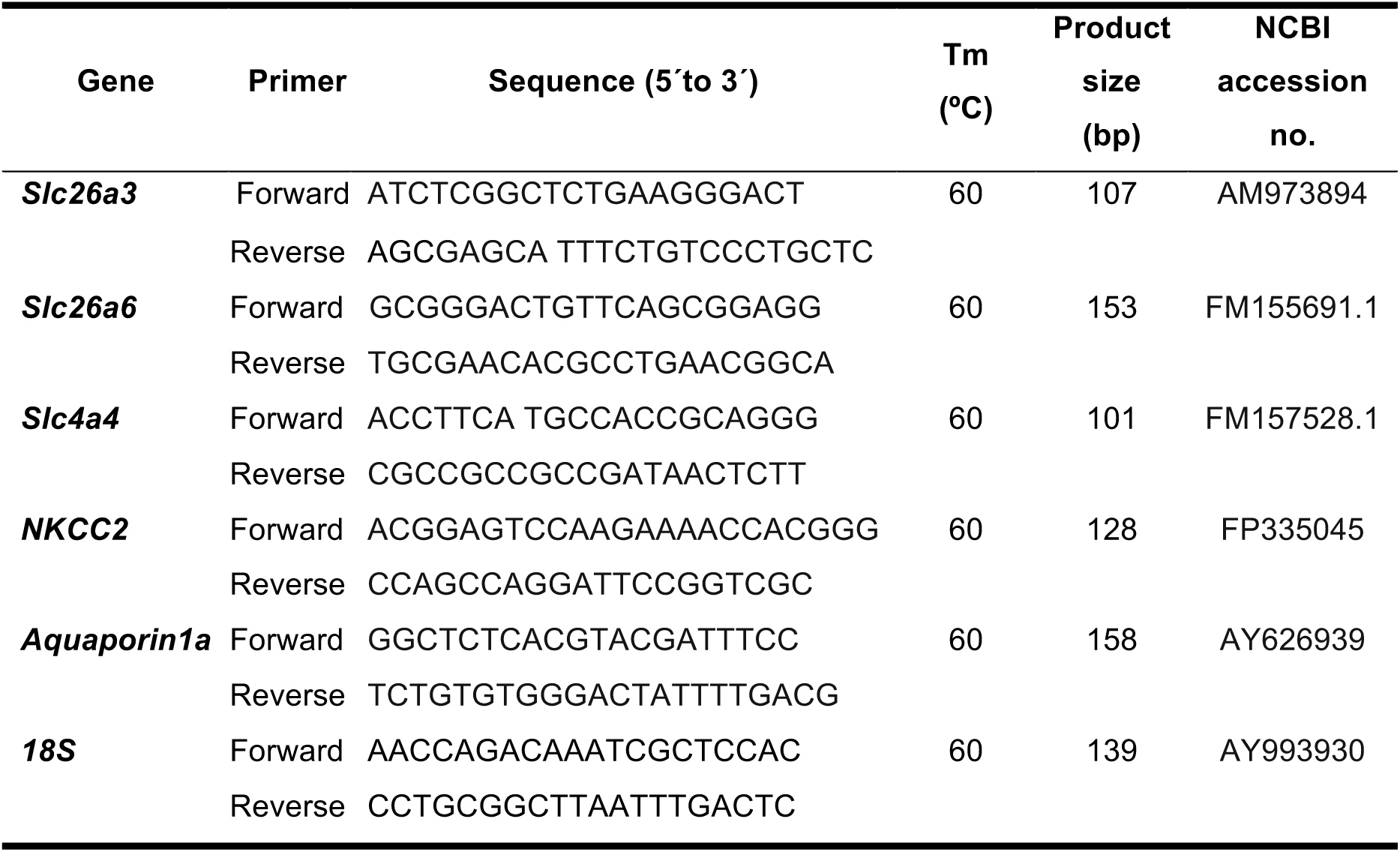
Details of primers used for qPCR.

**Figure 1.**
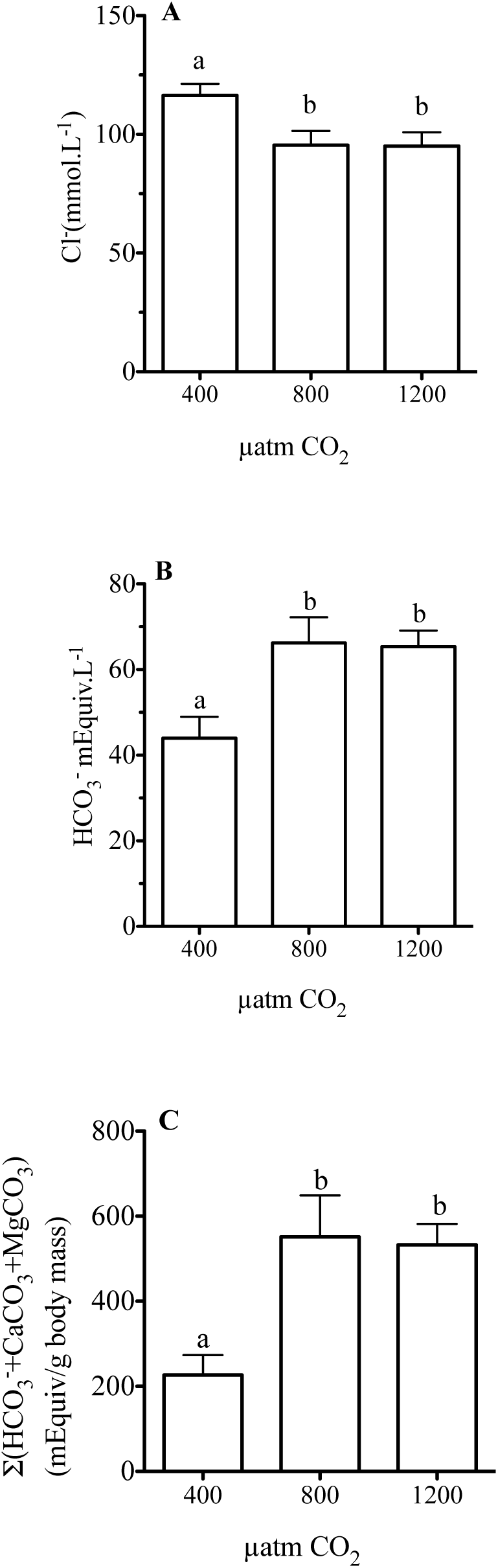
Characterization of Cl^−^, HCO_3_^−^ content in intestinal fluid and Ca(Mg)CO_3_ aggregates and total carbonate in intestinal lumen of sea bream in response to different environmental CO_2_ levels (400, 800 and 1200 μatm CO_2_) for up to three months. Each bar represents the mean ± s.e.m. (N= 7 - 9). Different superscript letters indicate significant differences (*p*<0.05, one-way ANOVA followed by Bonferroni *post hoc* test).

### Bicarbonate and carbonate aggregates in the intestinal lumen

HCO_3_^−^ concentration in the intestinal fluid of control fish was in the range of 44 mEq L^−1^ (Figure 1B) and the exposure to both levels of elevated CO_2_ resulted in a significant (*p*<0.05, one-way ANOVA) increase of HCO_3_^−^ concentration in the fish intestinal fluid to 66 and 65 mEq L^−1^ respectively (Figure 1B). When bicarbonate content was normalized to fluid volume plus the weight of carbonate aggregates (Ca[Mg]CO_3_) and fish body mass, we still observed that fish exposed to both levels of elevated CO_2_ responded with a 3-fold increase in carbonate aggregate formation (*p*<0.05, one-way ANOVA, Figure 1C).

### Bicarbonate secretion

HCO_3_^−^ secretion in the anterior intestine is shown in Figure 2A. In control fish (400 μatm CO_2_) bicarbonate secretion was 511 ± 7.3 nmol.h^−1^.cm^−2^, while it averaged 631 ± 7.2 and 842 ± 6.9 nmol.h^−1^.cm^−2^ for fish acclimated to 800 and 1200 μatm CO_2_, respectively. No significant effects were observed in bicarbonate secretion between the 400 and 800 μatm CO_2_ groups, while the 1200 μatm CO_2_-group was significantly higher than the control fish. In terms of epithelial resistance, no changes were observed between groups (Figure 2B).

**Figure 2.**
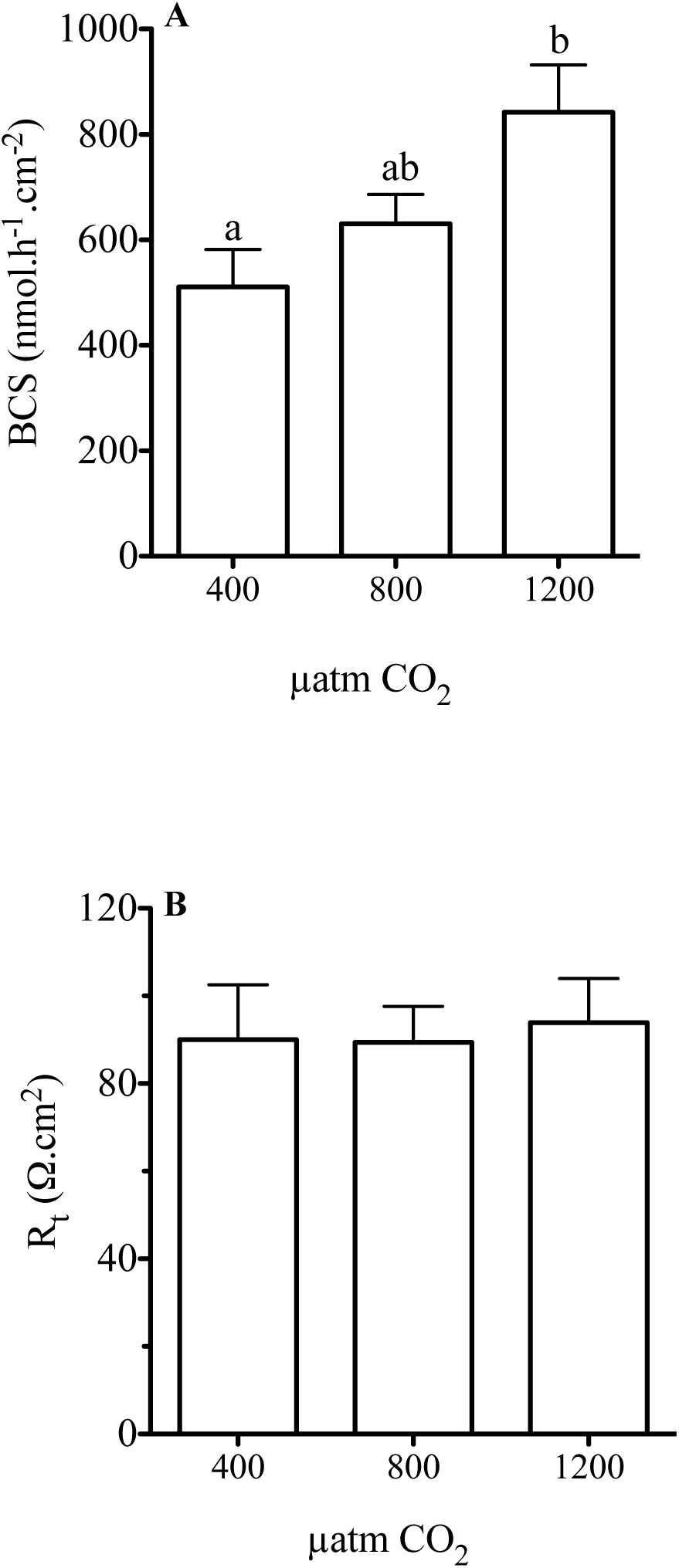
Bicarbonate secretion (BCS) and tissue resistance (Rt) measured in Ussing chambers by pH-Stat from anterior intestinal regions of sea bream in response to different environmental CO_2_ concentrations (400, 800 and 1200 μatm CO_2_) for up to three months. Each bar represents the mean ± s.e.m (N= 5 - 7). Different superscript letters indicate significant differences (*p*<0.05, one-way ANOVA followed by Bonferroni *post hoc* test).

### qPCR

The *Slc26a3* anion exchanger paralleled the accumulation of carbonate aggregates with a significant gradual increase of *Slc26a3* expression in response to increased water CO_2_ (Figure 3A). The apical exchanger *Slc26a6* presents a significantly (p<0.05, one-way ANOVA) higher expression in fish exposed to high water CO_2_ levels compared to controls (Figure 3B).

**Figure 3.**
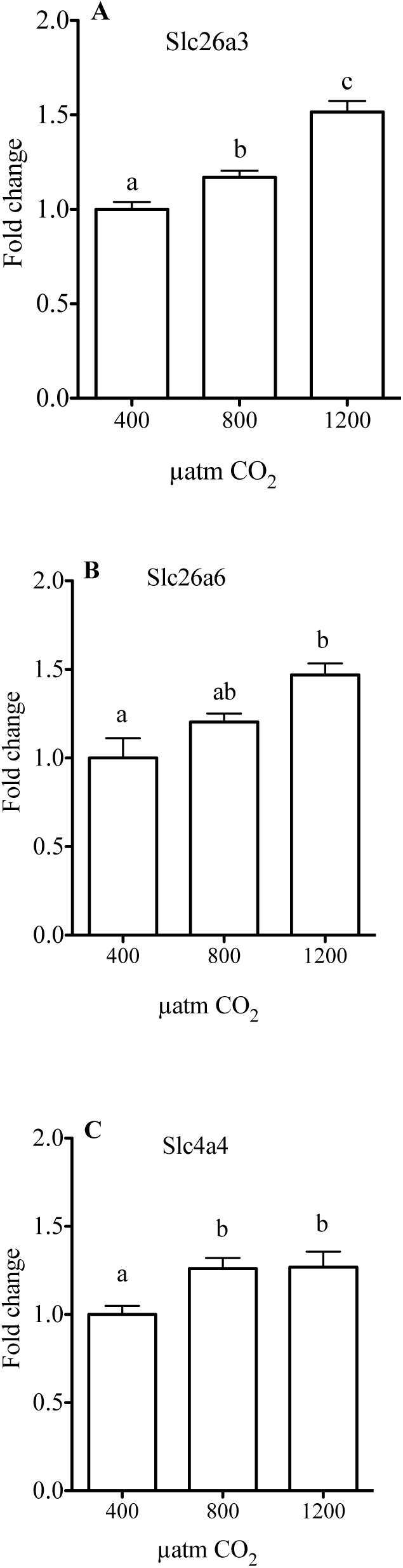
Relative expression (fold change of gene expression using *18S* as the housekeeping gene) of *Slc4a4, Slc26a3, Slc26a6* in the anterior intestine of sea bream in response to different environmental CO_2_ concentrations (400, 800 and 1200 μatm CO_2_) for up to three months. Each bar represents the mean ± s.e.m (N= 9 - 10). Different superscript letters indicate significant differences (*p*<0.05, one-way ANOVA followed by Bonferroni *post hoc* test).

Changes in *SLC4a4* expression are shown in Fig. 3C. Expression of the basolateral co-transporter *SLC4a4* was significantly (p<0.05, one-way ANOVA) higher in fish exposed to medium and high CO_2_ levels than in control fish (Figure 3E).

The expression of *NKCC2*, the absorptive form of the co-transporter in the anterior intestine of sea bream was significantly increased in response to hypercapnia (Figure 4A)

**Figure 4.**
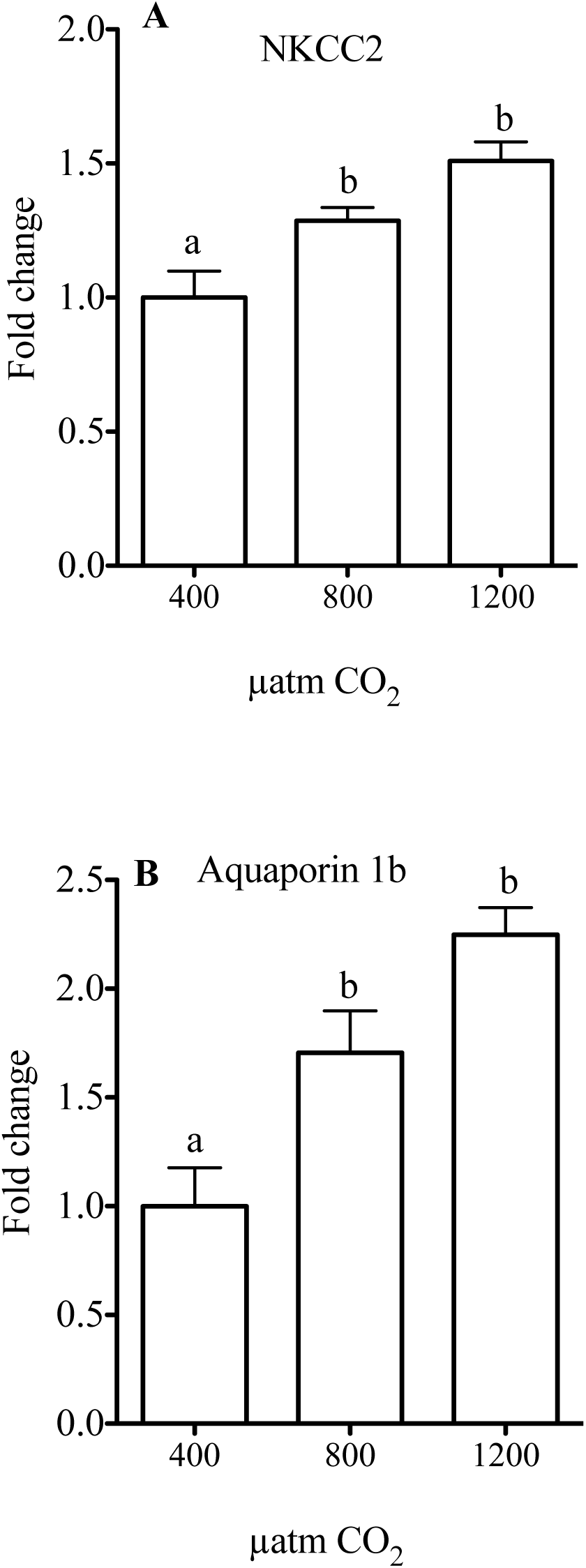
Comparative analysis of fold change of gene expression in sea bream juveniles using qPCR of *NKCC2* and *Aquaporin 1b* in the anterior intestine of sea bream in response to different CO_2_ concentrations (400, 800 and 1200 μatm CO_2_) for up to three months. Each bar represents the mean ± s.e.m (N= 8 - 11). Different superscript letters indicate significant differences (*p*<0.05, one-way ANOVA followed by Bonferroni *post hoc* test).

As expected, the water channel *Aquaporin 1b* also responded positively to the CO_2_ challenge, with significantly higher expression of *Aquaporin 1b* gene in fish exposed to medium and high CO_2_ concentrations than in controls (Figure 4B).

## DISCUSSION

The sea bream, acclimated from 400 up to 1200 μatm CO_2_ for three months, is able to maintain within a narrow range its plasma levels of osmolality, Na^+^ and Cl^−^ without suffering any casualties. This homeostatic balance is partially achieved by changes in intestinal epithelia related to ion transport. In this regard, elevated water CO_2_ induces increased rates of epithelial HCO_3_^−^ secretion and a molecular response that is in keeping with enhanced water absorption processes.

Gilthead sea bream acclimated to elevated water CO_2_ for 3 months increased the amount of HCO_3_^−^ and carbonate aggregates in the intestinal fluid (Figure 1). More precisely the groups of fish acclimated to hypercapnia (800 and 1200 μatm CO_2_), when compared to the control group (400 μatm CO_2_), showed a 1.5-fold and 3.0-fold increase in the bicarbonate and carbonate aggregates present in the intestinal lumen, respectively. These results and the fold-change of variation are in good agreement, with those previously described by us in seawater-acclimated sea bream challenged with 55ppt salinity (Gregorio et al., 2013). These effects in relation to salinity challenge, were related to an increase in water processing at the level of enterocytes. An enrichment of the intestinal fluid with Mg^2+^ and other divalent ions was described reflecting high levels of water ingestion and processing in the intestine (Kurita et al., 2008; Whittamore, 2012). Here, a noticeable increase of Mg^2+^ the intestinal fluid in response to hypercapnia is exposed, which suggests different water processing and/or drinking rates in response to higher water CO_2_.

In the intestinal fluid, both levels of hypercapnia elicited significant decreases in the intestinal fluid Cl^−^ content of about 21 mM. Interestingly, this decrease was paralleled by a ~22 mM increase in HCO_3_^−^ content of the same fluid, pointing to an activation of Cl^−^/ HCO_3_^−^ exchangers at the epithelial level (Grosell, 2006). To test this possibility, we measured bicarbonate secretion in the anterior intestine, since this portion of the intestine shows the highest secretion rates along the intestinal tract in the sea bream (Carvalho et al., 2012; Gregorio et al., 2013). Accordingly, intestinal bicarbonate secretion increased in parallel with increased environmental CO_2_ levels (Figure 2). The measured bicarbonate secretion fold-change between the control group and fish challenged with 1200 μatm CO_2_ from 511 to 842 nmol HCO_3_^−^ cm^−2^ h^−1^, coincided well with the alterations reported in the intestine of the sea bream challenged long term with 55 ppt seawater, from 495 to 783 nmol HCO_3_^−^ cm^−2^ h^−1^ (Gregorio et al., 2013). In this sense, the effects of hypercapnia could be carefully compared to a hyperosmotic challenge in this species, as they both enhanced bicarbonate secretion and formation of carbonate aggregates in the intestine.

It was previously reported that the seabream responds to hypercapnia with a plasma pH drop, which is buffered within the first 5 days of exposure by increasing plasma bicarbonate levels (Michaelidis et al., 2007). We hereby postulated that the accumulation of plasma bicarbonate generated in response to high environmental CO_2_, could be levelled off by activation of intestinal bicarbonate secretion at the level of the enterocytes. Plasma bicarbonate enters the enterocyte through an electrogenic Na^+^-coupled HCO_3_^−^ transporter located at the basolateral membrane, Slc4a4, which is important for a transepithelial HCO_3_^−^ transport (Kurita et al., 2008; Taylor et al., 2010). Our results show that the expression of *Slc4a4* increased in the groups challenged with high CO_2_ levels, supporting a hypothesis for enhanced bicarbonate secretion capacity in the intestine, in response to water hypercapnia and the consequent plasma HCO_3_^−^ accumulation.

In the apical membrane of the fish enterocytes, both bicarbonate secretion and chloride uptake are mediated through Slc26 family transporters, such as the Slc26a3 and Slc26a6 (Grosell et al., 2005; Kurita et al., 2008; Wilson et al., 1996). Likewise those exchangers are expressed in the intestine of sea bream and are modulated by salinity and endocrine regulation (Gregorio et al., 2014; Gregorio et al., 2013). In addition, the expression of this transporter is modulated in response to a CO_2_ challenge in the gill of toadfish (Esbaugh et al., 2012). Here, we confirmed that the expression of *Slc26a6* and *Slc26a3* are also modulated positively in response to elevated water CO_2_ levels in the intestine of the sea bream (Figure 3). This increase is in good agreement with the parallel expression increase of the *Slc4a4* co-transporter, which would substantiate a boosted secretory flow of bicarbonate from the plasma to the intestinal lumen, subsequently enhancing chloride absorption. It is challenging to establish transporters stoichiometry from ion concentration alone. However, here we have established a 1:1 relationship between the variation of Cl^−^ and HCO_3_^−^ in the intestinal fluid after 3 months in hypercapnia (Figures 1A and B). It appears, that as a consequence of the increased bicarbonate levels on the intestinal fluid, carbonate aggregate formation is increased, sustaining a decrease of fluid osmolality and that enables water absorption (Wilson et al., 2002).

In marine fish, water absorption in the intestinal tract is mediated through the activity of ion absorption, especially Cl^−^. One of the most important ion transporters in this species is the Na^+^-K^+^-2Cl^−^ co-transporter (NKCC) (Musch et al., 1982), which mediates the electroneutral movement of 1 Na^+^, 1 K^+^ and 2 Cl^−^ across cell membranes. Two isoforms, *NKCC1* and *NKCC2,* are currently known and are encoded by different genes (Flatman, 2002). In the intestine of marine fish the apical co-transporter NKCC2 activity seems essential for ion regulation and, in parallel with the Slc26a6, it is believed to regulate water homeostasis in salinity-challenged sea bream (Gregorio et al., 2013; Grosell, 2006). Here, we show that elevated water CO_2_ causes an upregulation of *NKCC2* expression in the anterior intestine (Figure 4A) which adds up to increased expression of the apical anion transporters, *Slc26a6* and *Slc26a3.* This combination has the potential to enhance intestinal water absorption linked to chloride movements. Remarkably, sea bream exposed to elevated water CO_2_ showed no significant changes in plasma ions concentration, indicating that the intestinal response to elevated water CO_2_, is part of the allostatic control of plasma homeostasis. In order to understand these changes further, we analysed the expression of *Aquaporin 1b,* a functional water channel highly expressed in intestinal epithelial cells of the sea bream anterior intestine (Martos-Sitcha et al., 2015; Raldua et al., 2008). We observed solid increases of *Aquaporin 1b* expression in the anterior intestine of sea bream exposed to elevated water CO_2_. Providing further evidence of the functional and molecular changes of the intestine in response to elevated water CO_2_, that could be the foundation of increased energetic expenditure recently reported in the toadfish intestine in response to elevated water CO_2_ (Heuer and Grosell, 2016).

## ACKNOWLEDGEMENTS

We appreciate the technical input of Dr. João Eugenio Reis (Ramalhete Marine Station, CCMar, University of Algarve, Portugal) for fish care and assistance during experimental periods.

